# Natural body-selective motion processing in humans: Evidence from event-related potentials

**DOI:** 10.1101/2025.01.08.631855

**Authors:** Jane Chesley, Lars Riecke, Beatrice de Gelder

**Author notes:** Correspondence to: Room 3.009, Oxfordlaan 55, 6229 EV Maastricht, the Netherlands. E-mail address (B. de Gelder).

## Abstract

As social species, perception of body motion is crucial to daily interactions, yet the underlying neural processes and their dynamics are not fully understood. While well-established EEG evidence has identified temporal markers of biological body motion, and recent fMRI research points to naturalistic body networks, the temporal markers of higher-level, naturalistic body motion processes remain unknown. The present study bridges this gap, showing that at the stage of P2, the cortex may already distinguish between the global motion of naturalistic bodies compared to objects. Exploratory analyses point to a similar body-specific modulation across stimulus species, namely human and monkey. The findings of the present study expand our understanding of body motion processes, in line with emerging, integrative models of multi-faceted body processes.

## 1. Introduction

Body motion is crucial to social interactions in both humans and non-human primates. As social species, we perceive bodily cues from our conspecifics, and we inherently use that information to inform our daily trajectory. Yet, the neural dynamics of the underlying body motion processes are not fully understood.

From a traditional perspective, body processes are thought to occur in the extrastriate body area (EBA) and fusiform body area (FBA), with functional selectivity that is distinct from that of face or object processes (Downing et al., 2001; Peelen & Downing, 2005, 2007; Pourtois et al., 2007; Schwarzlose et al., 2005). However, further evidence has accumulated for body-selectivity widespread beyond EBA and FBA, including the superior temporal sulcus (Candidi et al., 2015; Kret et al., 2011; Pichon et al., 2012), temporoparietal junction (TPJ), premotor cortex (Pichon et al., 2009), and subcortical areas (for reviews see de Gelder, 2006; de Gelder & Poyo Solanas, 2021).

On a similar note, traditional temporal investigations of body processes have localized a body-selective event-related potential (ERP) which occurs 170-190 ms post-stimulus (N1) and is apparent when comparing body images to their scrambled counterparts (Gliga & Dehaene-Lambertz, 2005), faces (Thierry et al., 2006), plants (Moreau et al., 2018; Taylor et al., 2010) and inversions (Stekelenburg & de Gelder, 2004). This body-specific N1 modulation is well-established and has been further corroborated by intracranial recordings directly from EBA, showing body-selective responses 190 ms after body image onset (Pourtois et al., 2007).

However, a crucial feature of bodies is their physical dynamics: As social emotional expressions, or intentional actions. Yet, despite the importance of the integration of body and motion, most of what is known about body processing is based on experimental designs employing motionless body images, as in the aforementioned studies, rather than bodies in motion.

Recent fMRI research corroborates this integrative perspective, as body-selective processes have been shown to overlap with the neural processing of biological movement (Jastorff & Orban, 2009), postural features (Marrazzo et al., 2023; Marrazzo et al., 2021; Poyo Solanas et al., 2020), action recognition (Goldberg et al., 2014; Shmuelof & Zohary, 2005), and social perception (Kret et al., 2011; Moreau et al., 2023). Most notably, emerging evidence points to a large-scale neural network specifically selective for naturalistic body motion in the human brain (Li et al., 2023) as well as the non-human primate brain (Kumar et al., 2023).

In the temporal domain, body motion processes have mainly been investigated with point-light-display (PLD) stimuli, which present non-naturalistic, moving dots to mimic biological body motion (Johansson, 1973). Several PLD studies in humans have found clear differences between biological and scramble motion, which are apparent in negative ERP components such as N1 and N2 (Hirai et al., 2003, 2005; Jokisch et al., 2005; White et al., 2014), as well as positive components such as P1 and P2 (Buzzell, 2013; Pegna, 2015; Krakowski et al., 2011). Further ERP research suggests that as young as eight months, the human brain may already discriminate biological PLD motion (Reid et al., 2008).

The findings of PLD studies suggest there are clear temporal differences in the processes of biological versus non-biological body motion. However, due to the are also true for naturalistic representations of body motion. In addition, it is still unclear whether the body motion effects observed in PLD studies are truly body-specific or rather generalized to motion effects for other animate stimuli, such as faces.

The present study aims to address these gaps by investigating the temporal patterns of naturalistic, body-specific motion processing. We employed electroencephalography (EEG) and measured ERPs to assess the millisecond-precise timing of body motion processing. We presented video clips of natural body motion, exclusive of face information, to human subjects. To isolate higher-level body processes, we included scrambled versions of the video clips that control for the contributions of low-level visual features, such as luminance, contrast and non-background area. To target body-selective responses, we experimentally manipulated the category of the moving stimulus across three conditions: body, face, object. Furthermore, to allow exploring whether the putative body-selectivity was specific to the bodies of humans, we varied the species of the animate stimuli, by presenting video clips of humans as well as monkeys.

We hypothesized that event-related cortical activity within the P1-N2 range would be modulated by category, with a naturalistic body motion-specific response over temporo-parieto-occipital scalp regions which were previously shown to be modulated by body point-light motion (Krakowski et al., 2011). In addition, we expected that this putative effect would be observable with human motion stimuli as well as monkey motion stimuli.

## 2. Methods

### 2.1 Ethics statement

Procedures were approved by the Ethical Committee of Maastricht University and were in accordance with the Declaration of Helsinki. Written, informed consent was obtained from participants prior to the experiment, and the study was conducted in accordance with local legislation and institutional requirements.

### 2.2 Participants

Thirty healthy, right-handed participants with normal or corrected-to-normal vision were recruited for this study. All participants reported no history of psychiatric or neurological disorders. Participants were compensated in either monetary vouchers (€7.5 per hour) or credit points (1 credit per hour). One participant’s data were excluded from the analysis because she/he reportedly confused the response buttons in the attention task (as shown by 0% accuracy); the remaining 29 participants had an average accuracy of 96 ± 5% (mean ± SD) (range = 81 – 100%). Hence, 29 participants’ data were included in the analysis (17 females; age range = 18-37 years; mean age = 23).

### 2.3 Stimuli

Grayscale, naturalistic videos of bodies, faces and objects were used as stimuli in the experiment. Body and face stimuli were from a human or a monkey. Body stimuli had face information removed with Gaussian blurring. Object stimuli consisted of two sets of artificial objects (e.g., mechanical devices, vehicles, tools) and their aspect ratio matched either human bodies (set 1) or monkey bodies (set 2). Stimuli were embedded in a white noise background and presented centrally on the computer 20 degrees for human bodies and objects, and 16 * 16 degrees for monkey faces, bodies, and objects.

To control for the contribution of low-level visual features, mosaic-scrambled videos were included. Mosaic-scrambled videos destroyed the whole shape of each body/face/object stimulus but preserved the low-level features of luminance, contrast, and non-background area (Bognár et al., 2023). This resulted in a total of six experimental conditions (body/face/object * normal/scrambled). There were twenty different stimuli per condition, with an equal distribution of human and monkey actors, resulting in 120 unique videos.

All videos were used in previous studies on body and face processing (Li et al., 2023; Bognár et al., 2023; Kret et al., 2011; Zhu et al., 2013). The videos were 1-second in duration, presented at 60 frames per second. The body videos depicted either a human or a monkey performing naturalistic full-body movements, and the face videos depicted either a human or a monkey performing naturalistic facial movements. The human body and human face videos depicted both female and male actors dressed in black, performing expressions against a greenscreen background (Kret et al., 2011). The expressions included full-body or facial expressions of anger, fear, happiness, as well as neutral actions such as nose-pulling or coughing. The original monkey videos were recorded from rhesus monkeys from the Katholieke Universiteit Leuven monkey colony. The monkey body videos depicted full-body movements such as grasping, picking, turning, walking, threatening, throwing, wiping, and initiating jumping (Bognár et al., 2023). The monkey face videos depicted facial expressions such as chewing, lip-smacking, fear both emotional and neutral poses were included. The object videos depicted non-rigid movements of computer-rendered artificial objects (created by https://garethwashere.tumblr.com) (Bognár et al., 2023).

Stimulus presentation was programmed in MATLAB 2021a (The Mathworks, Natick, MA, USA) with the Psychophysics Toolbox extensions (Brainard, 1997; Pelli, 1997; Kleiner et al., 2007) as well as custom code.

**Figure 1.**
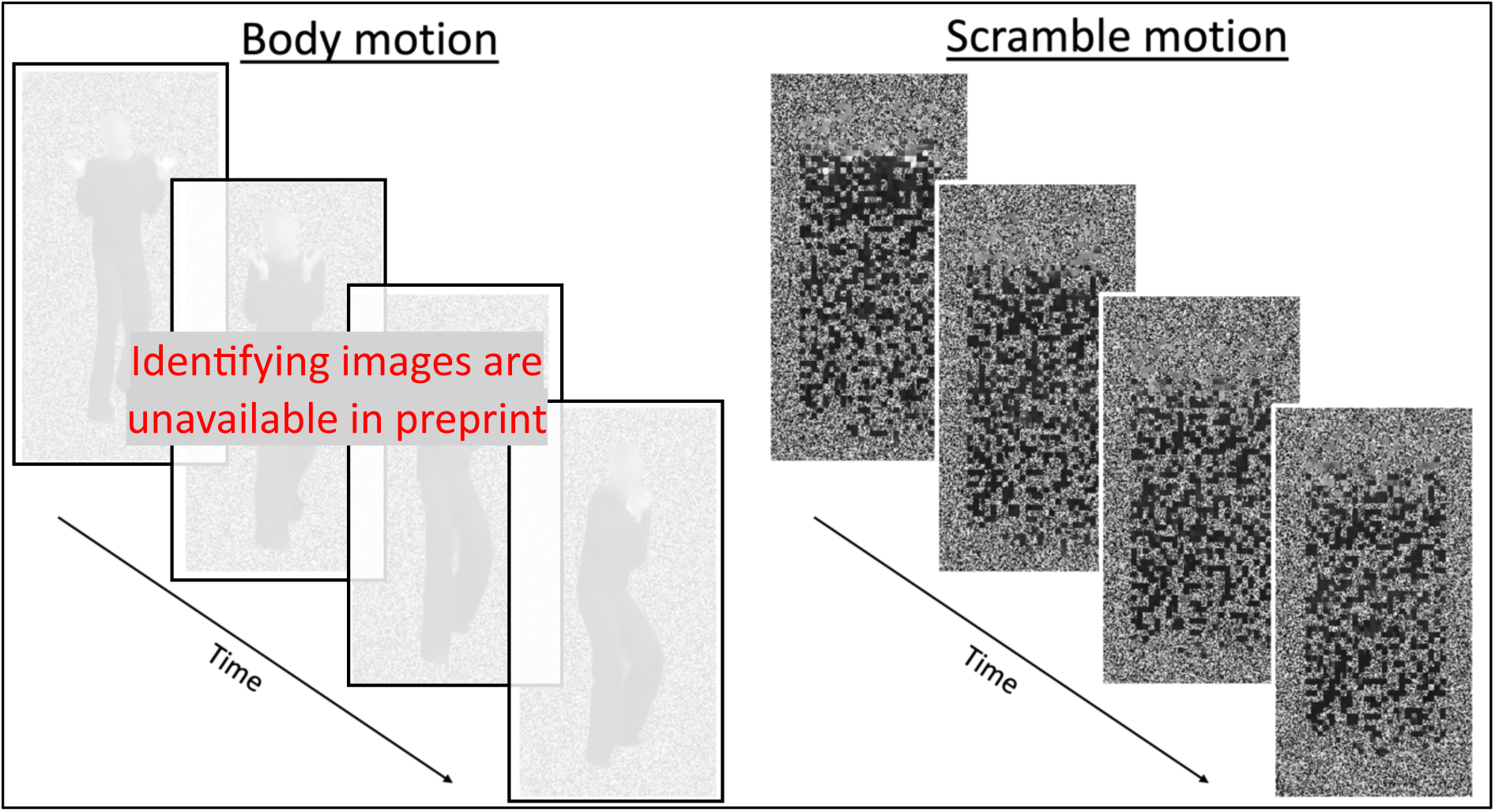
Example stimuli. Image sequences are shown for the body motion condition (left) and scramble body motion condition (right). For visualization purposes, four frames are shown for each stimulus. In practice, each stimulus was presented as a video (1s) at 60 frames per second, with a white fixation cross centered and overlaid on each video.

### 2.4 Experimental design, task and procedure

The experimental procedure consisted of two sessions, one of which presented videos and the second of which presented static images (a single frame) extracted of the same video stimuli. The order of the two experimental sessions was randomized across participants. The present paper reports the methods, analysis, and results of the former, video-related experimental session; the latter was reported in a previous study (Chesley et al., 2024).

The main experiment employed a fully crossed within-subject factorial design. There were four runs of the experiment, all lasting around 6 minutes. During each run, 120 unique videos (6 conditions × 20 stimuli; see Stimuli) were presented once in random order. This resulted in a total of four repetitions per stimulus and 80 repetitions per condition. Each trial began with a white fixation cross centered on a gray screen, which remained visible throughout the trial. To reduce the temporal expectancy of stimulus presentation, the intertrial interval was jittered at 1500 ms (1500 ± 200 ms). Participants viewed the videos on a computer screen (1920 × 1080) at 65 cm from their eyes. Participants were asked to focus their gaze on the fixation cross and focus their attention on each stimulus. To maintain attention, a question appeared on a random 10% of trials. The question asked about the content of the preceding stimulus (E.g., “What did the previous video show?”), and participants were asked to respond with a button press from a selection of “Body”, “Face”, “Object” or “None of the above.”

### 2.5 EEG acquisition

EEG signals were acquired from 33 passive silver/silver chloride electrodes embedded in a fabric cap (EASYCAP GmbH) and arranged in accordance with the international 10-20 system. Scalp electrodes included: AFz, Fz, FCz, Cz, CPz, Pz, Oz, Fp1, Fp2, F3, F4, F7, F8, FC3, FC4, FT7, FT8, C3, C4, T7, T8, CP3, CP4, TP7, TP8, TP9, TP10, P3, P4, P7, P8, O1, and O2 (n = 33). EEG signals were amplified with a BrainVision amplifier (Brain Product GmbH, Germany) and recorded with BrainVision Recorder (Brain Product GmbH, Germany) at a sampling rate of 1000 Hz. Electrooculogram signals were recorded from two electrodes placed 1 cm from the outer canthi of each eye, as well as from two electrodes placed 1 cm above and below the center of one eye. An online reference electrode was placed on the left mastoid and an offline reference electrode was placed on the right mastoid. The ground electrode was placed on the forehead. Impedance was kept below 5 kΩ for all electrodes. EEG recordings took place in an electromagnetically shielded room.

### 2.6 EEG data preprocessing

EEG data were preprocessed and analyzed offline in MATLAB 2021a (The Mathworks, Natick, MA, USA) using the Fieldtrip Toolbox extensions (Oostenveld et al., 2011) as well as custom code. The signal was first segmented into trials from 600 ms pre-stimulus onset (video presentation) to 1700 ms post-stimulus. EEG data were re-referenced to the average of the signal recorded at the left and right mastoids and downsampled to 250 Hz. Ocular movements were removed with Independent Component Analysis (ICA, logistic infomax ICA algorithm; Bell & Sejnowski, 1995); on average, 2.3 ± 0.7 (mean ± SD) (range = 1 – 4) eye movement-related components were visually identified and removed per participant. The channel signals were filtered with a 0.3 – 20 Hz bandpass filter. Single trials in which the peak amplitude exceeded +/− 75 mV were rejected; on average, 2.1 ± 5.8% (mean ± SD) (range = 0 – 31.7%) of trials were rejected per participant. Baseline correction was performed by subtracting the mean signal during the pre-stimulus period (−50 to 0 ms) from each time point in the epoch of interest (0 – 1000 ms post-stimulus).

### 2.7 ERP analysis

#### 2.7.1 Region-of-interest and ERP Components

The pre-processed signal was averaged across six channels (F7, TP7, PO7, F8, TP8, and PO8) within the scalp region of interest (ROI). This ROI was selected based on previous literature suggesting body motion differs from scramble motion at bilateral temporo-parieto-occipital channels (see Introduction).

Based on previous research on body motion processing, components P1, N1, P2 and N2 were of interest (see Introduction). To measure these components in the present data, average waveforms from the ROI were collapsed across all conditions (Fig. 2) and peak amplitudes were quantified as the maximum positivity or negativity, respectively for positive and negative components. Component time-windows were defined by centering 20-30 ms time-windows around each peak amplitude. This resulted in three component time-windows, namely N1: 174 +/− 15 ms, P2: 218 +/− 15 ms, and N2: 262 +/− 10 ms. Component P1 was not observed in the present data, which may be attributed to the dynamic and variable nature of the stimuli, which started immediately from their onset.

**Figure 2.**
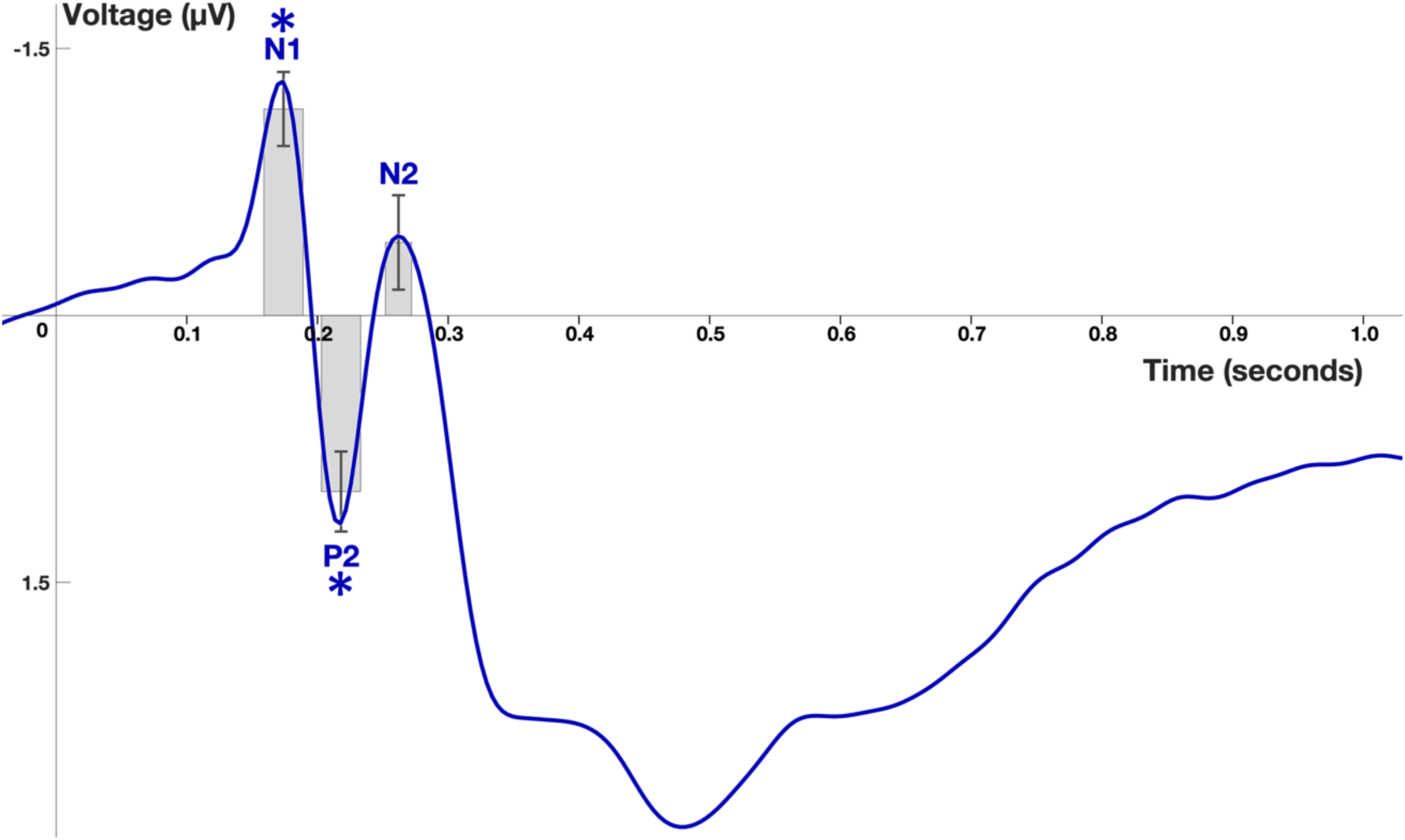
Grand-averaged ERP across all conditions, extracted from bilateral temporo-parieto-occipital channels. Three components were observed: N1 (174 +/− 15 ms), P2 (218 +/− 15 ms) and N2 (262 +/− 10 ms). Gray boxes represent component time-windows (x-range) as well as the mean voltage across all subjects (y-range) with standard error bars. The mean voltage of components N1 and P2 were statistically different from zero (*p*-adj < 0.05), as indicated by asterisks. The mean voltage of component N2 was not statistically different from zero, although significance was approached (*p*-adj = 0.066).

In addition, to determine whether the observed components were significantly different from zero, three one-sample t-tests with False Discovery Rate (FDR) correction for multiple comparisons (Benjamini & Hochberg, 1995) were performed on the mean amplitude within each time-window.

#### 2.7.2 Differential waveforms

To control for potentially confounding effects of low-level visual features, the each category (Fig. 3). For a given waveform, the subject-level mean amplitude for each scramble category was subtracted from the respective mean amplitude for each normal category: Body (normal – scramble); Face (normal – scramble); Object (normal – scramble). The resulting differential waveforms were deemed to represent the ERP responses related to higher-level visual processes and were further analyzed. To further assess the significance of the higher-order differential ERPs, one-sample t-tests with FDR correction for multiple comparisons were performed on the mean differential amplitude for each component and category.

**Figure 3.**
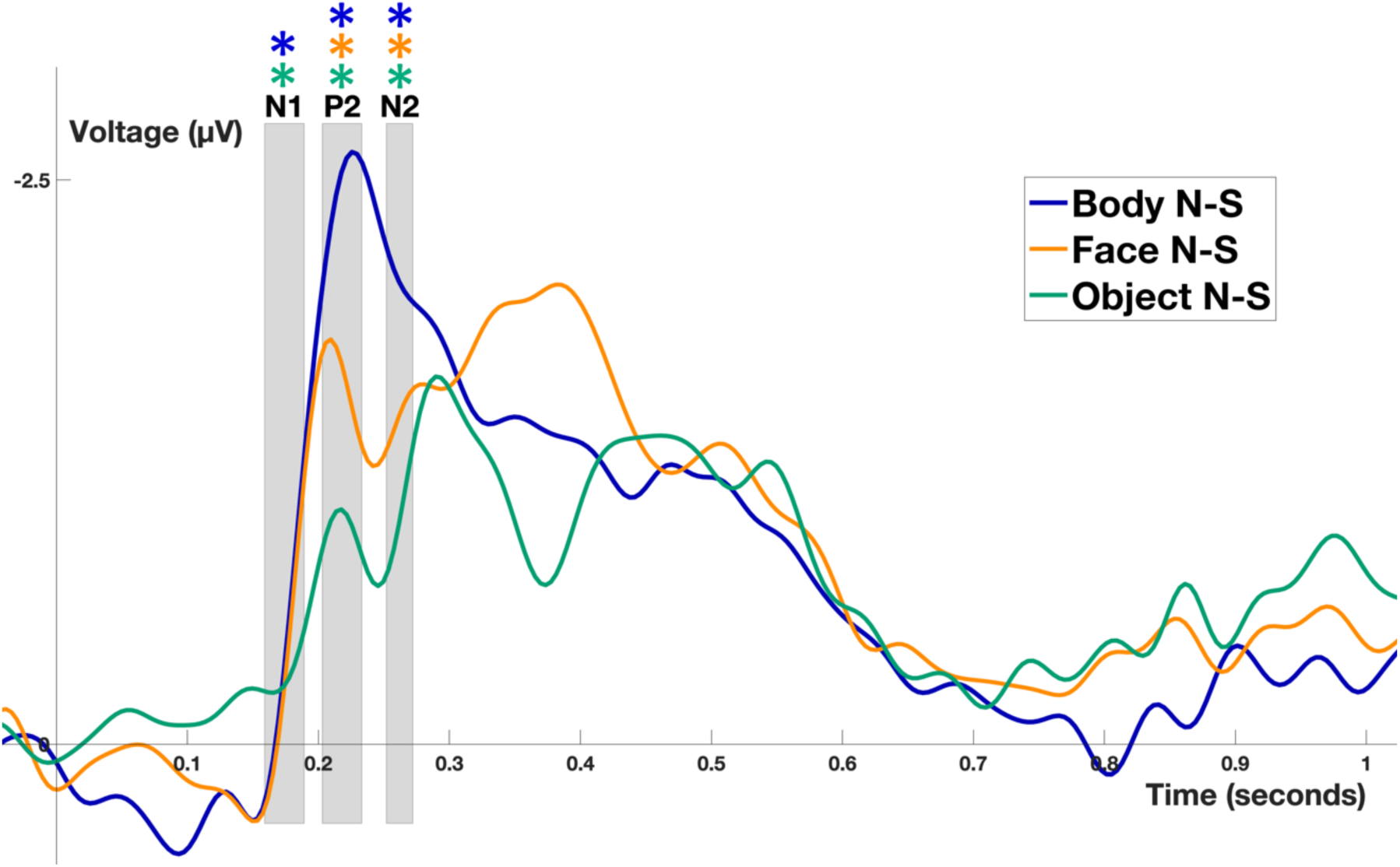
Differential ERPs (Normal – Scramble) for each category: Body (blue), Face (orange) and Object (green), averaged across all subjects. Gray boxes represent component time-windows. One-sample t-tests revealed the differential mean amplitudes for all categories during P2 and N2 components were significantly lower than zero (*p*-adj < 0.05), as indicated by asterisks. For component N1, the differential mean amplitudes for Body and Object categories were significantly lower than zero (*p*-adj < 0.05), but this was not significant for the Face category.

### 2.8 Statistical analyses

Statistical analyses were performed using R (Version 4.3.3; R Foundation for Statistical Computing, 2023) and RStudio (Version 2023.09.1+494; Posit Software, PBC, 2022). Before performing parametric statistical tests, all distributions were verified for normality and values exceeding the interval mean ±3 SD were considered as outliers and discarded. To assess the effect of category, a one-way ANOVA with three levels (Body/Face/Object) was applied to the mean differential amplitude (normal – scramble) for each component. FDR correction was applied to correct for multiple comparisons; only corrected p-values are reported. Given a significant effect of category, post-hoc pairwise comparisons were performed with Tukey’s HSD, which inherently corrects for multiple comparisons. Statistical differences below p < 0.05 were considered significant.

## 3. Results

### 3.1 ERP components

Results of one-sample t-tests showed the grand-averaged amplitude of component N1 (M = −1.16; SD = 1.12) was significantly lower than zero (*t*(28) = −5.55, *p*-adj < .001, one-tailed) and the grand-averaged amplitude of component P2 (M = 0.99; SD = 1.21) was significantly higher than zero (*t*(28) = −4.43, *p*-adj < .001, one-tailed) (Fig. 2). The grand-averaged amplitude of component N2 (M = −0.41; SD = 1.43) was not statistically lower than zero (*t*(28) = −1.55, *p*-adj = 0.066, one-tailed), although significance was approached.

### 3.2 Differential ERP components

Initial visual inspection of the differential (Normal – Scramble) ERP waveforms indicated a relatively long-lasting higher-order motion process may occur, as indicated by a strong and sustained negativity between ∼170 ms to ∼800 ms post-stimulus (Fig. 3). In addition, a strong and temporally concise peak is centered around the P2 time window, followed by a more sustained pattern extending beyond the N2 time window.

To test our hypothesis, the differential ERP amplitudes within the time windows of interest were statistically analyzed. Results of one-sample t-tests confirmed the observations above, showing higher-order differential ERPs were consistently different from zero (Fig. 3). Namely, for component N1, the mean differential amplitude of Body (M = −0.36; SD = 0.87; *t*(28) = −2.25, *p*-adj = 0.02, one-tailed) and Object (M = −0.29; SD = 0.74; *t*(28) = −2.09, *p*-adj = 0.03, one-tailed) categories was significantly lower than zero. The mean differential amplitude of Faces for component N1 was not statistically lower than zero (M = −0.29; SD = 0.99; *t*(28) = −1.56, *p*-adj = 0.07, one-tailed), although significance was approached. For component P2, the mean differential amplitude of Body (M = −2.26; SD = 1.22; *t*(27) = −9.8, *p*-adj < 0.001, one-tailed), Face (M = −1.64; SD = 1.16; *t*(28) = −7.64, *p*-adj < 0.001, one-tailed), and Object (M = −0.98; SD = 1.05; *t*(28) = −5.03, *p*-adj < 0.001, one-tailed) categories was significantly lower than zero. For component N2, the mean differential amplitude of Body (M = −2.06; SD = 1.16; *t* (28) = −9.56, *p*-adj < 0.001, one-tailed), Face (M = −1.45; SD = 1.38; *t*(28) = −5.66, *p*-adj < 0.001, one-tailed), and Object (M = −0.99; SD = 0.98; *t*(28) = −5.44, *p*-adj < 0.001, one-tailed) categories was significantly lower than zero.

### 3.3 Main results

Component P2 analysis revealed a significant effect of category on mean differential amplitude, *F*(2, 83) = 8.91, *p*-adj = 0.001, *η*^2^ = 0.18 (Fig. 4A). Post-hoc pairwise comparisons revealed this effect was driven by a significant difference between body and object categories, mean difference (MD) = 1.28, 95% CI [-0.56, 2.00], *p*-adj < 0.001. No significant difference was observed between face and object categories (MD = 0.66, 95% CI [-0.06 1.38], *p*-adj = 0.08), nor between face and body categories (MD = 0.62, 95% CI [-0.11 1.34], *p*-adj = 0.11).

**Figure 4.**
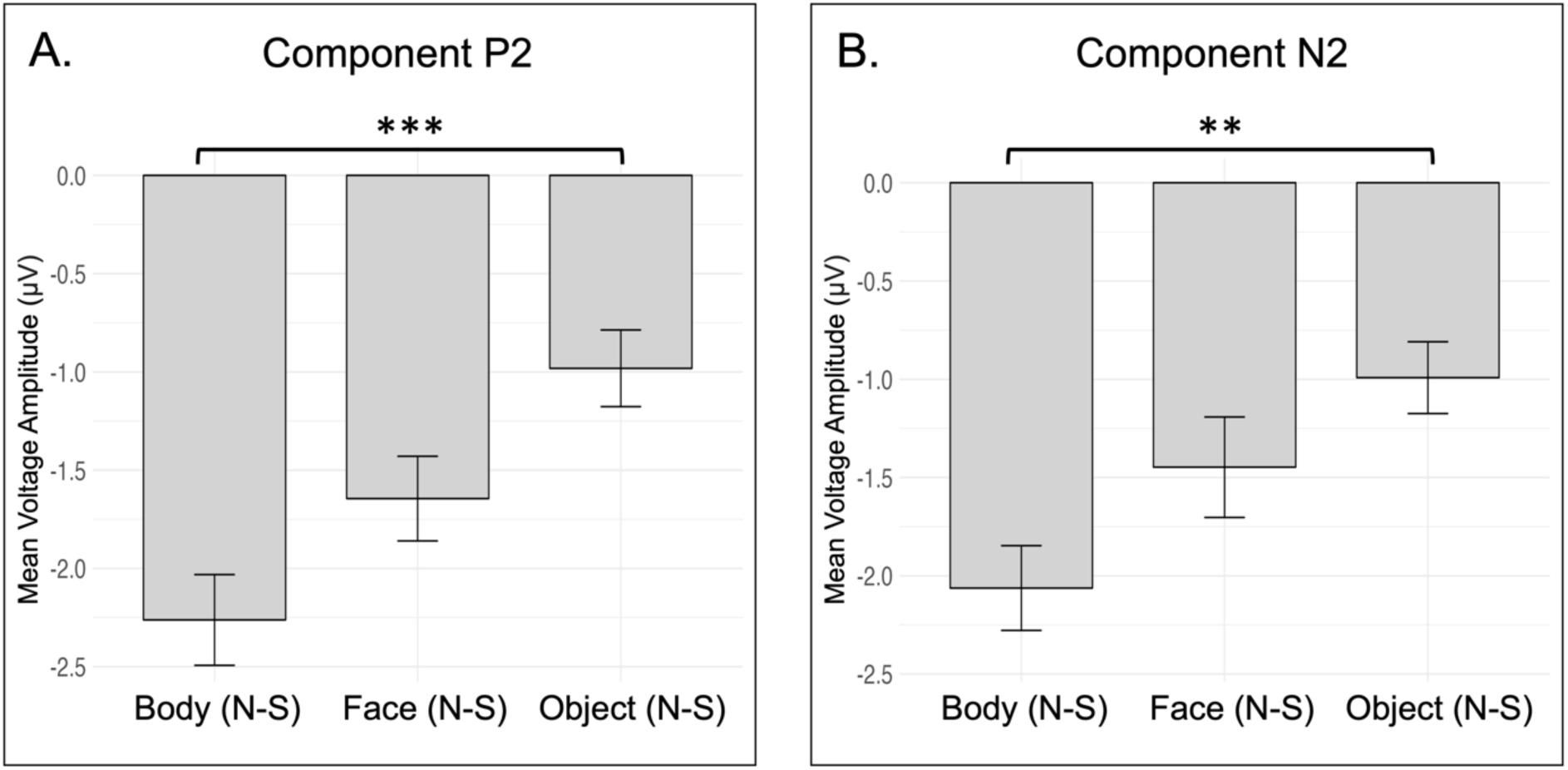
Means of differential voltage amplitude (Normal – Scramble) across categories, shown separately for component P2 (A) and component N2 (B). One-way ANOVAs revealed a significant category effect on components P2 and N2 (*p*-adj < 0.05). This category effect was not significant for component N1. Pairwise comparisons showed a statistically significant difference between body and object conditions for component P2 (*p*-adj < 0.001) and component N2 (*p*-adj = 0.003), as indicated by asterisks.

Similar to the P2 results above, component N2 analysis revealed a significant effect of category on mean differential amplitude, *F*(2, 84) = 5.96, *p*-adj = 0.006, *η*^2^ = 0.12 (Fig. 4B). Post-hoc pairwise comparisons revealed this effect was also driven by a significant difference between body and object categories, MD = 1.07, 95% CI [0.33, 1.81], *p*-adj = 0.003. Again, no significant difference was observed between face and object categories (MD = 0.46, 95% CI [-0.29 1.20], *p*-adj = 0.31), nor between face and body categories (MD = 0.61, 95% CI [-0.13 1.36], *p*-adj = 0.12).

Component N1 analysis revealed no significant effect of category on mean differential amplitude, *F*(2, 84) = 0.07, *p*-adj = 0.93, *η*^2^ = 0.002.

### 3.2 Post-hoc, exploratory analyses and results

Post-hoc, exploratory analyses were performed to further characterize the observed effect of category on differential voltage amplitude for component P2. Component N2 was not further investigated because this component was observed less reliably across participants: unlike the P2, the grand-averaged amplitude for component N2 did not differ significantly from zero (*p*-adj = 0.066) and the differential waveforms showed no demarcated peak during the N2 interval (see ERP analysis).

To better understand the categorical profile of the observed P2 effect, data were split by species. The original stimuli included videos of both human and monkey bodies and faces, as well as artificial objects with the aspect ratio matched to human bodies and monkey bodies (see Stimuli). Hereafter, artificial object categories with the aspect ratio matched to human bodies are referred to as ‘human object’, and artificial object categories with the aspect ratio matched to monkey bodies are referred to as ‘monkey object.’

To investigate whether the observed P2 modulation by category was species-specific, differential ERP waveforms (normal-scramble) were calculated separately for human conditions (human body, human face, and human object) as well as monkey conditions (monkey body, monkey face, and monkey object) (Fig. 5). Two one-way ANOVAs with three levels (Body/Face/Object) were applied to the mean differential voltage (normal – scramble) for component P2, respectively for human categories and monkey categories.

**Figure 5.**
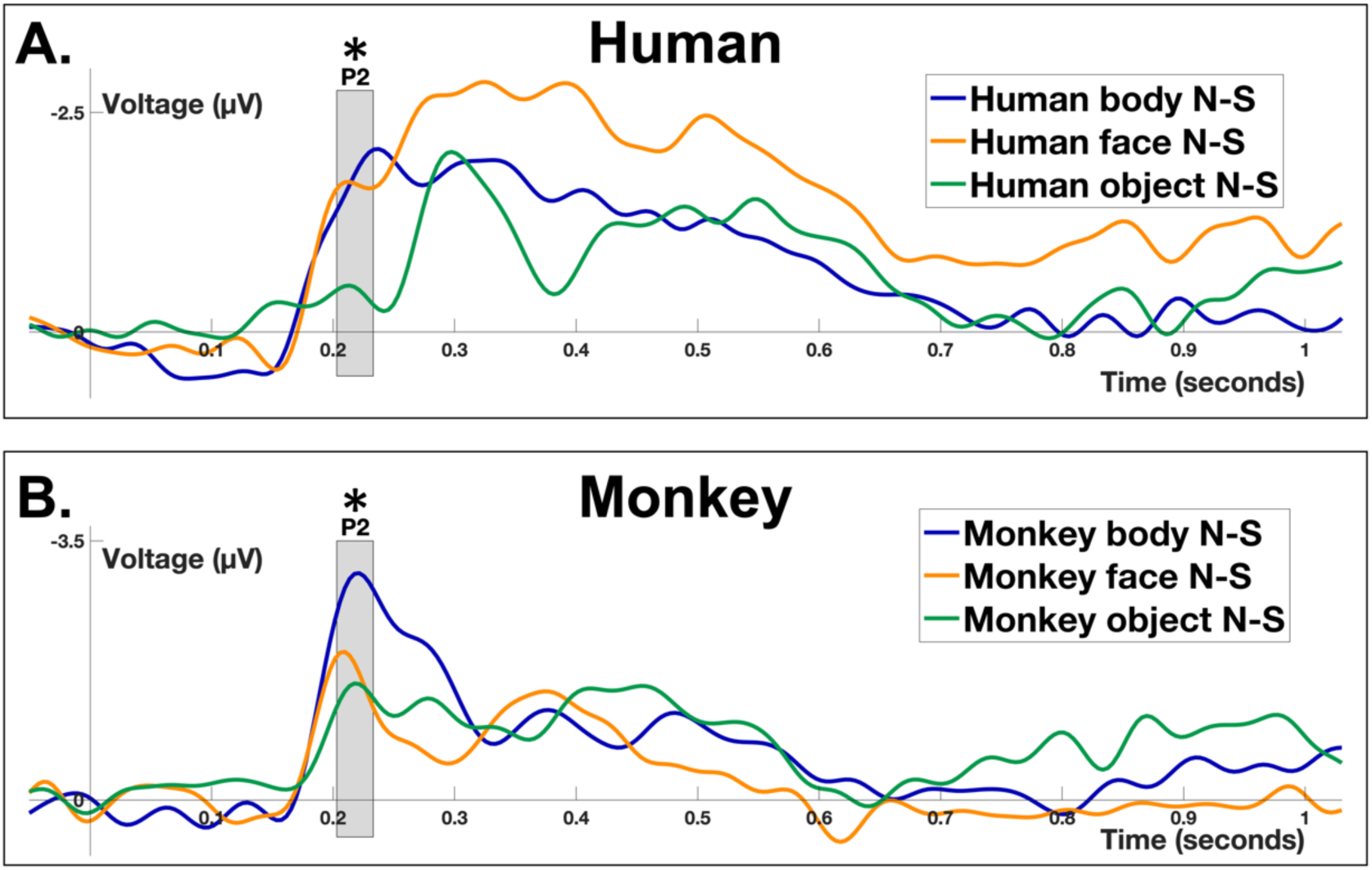
Differential ERPs (Normal – Scramble) for each category: Body, Face, and Object, shown separately for Human (A) and Monkey (B) categories. The grey box represents component P2 time-window, which was used for further analyses. One-way ANOVAs revealed significant P2 category effects among both human (*p*-adj = 0.001) and monkey (*p*-adj = 0.001) categories, as indicated by asterisks.

Analysis of human categories revealed a significant effect of category (*F*(2, 83) = 7.28, *p*-adj = 0.001, *η*^2^ = 0.15), driven by a significant difference between human body and human object categories (MD = 1.29, 95% CI [0.39, 2.20], *p*-adj = 95% CI [0.30, 2.10], *p*-adj = 0.006) (Fig. 6A). No significant difference was observed between human body and human face categories (MD = 0.09, 95% CI [-0.81, 1.00], *p*-adj = 0.967).

**Figure 6.**
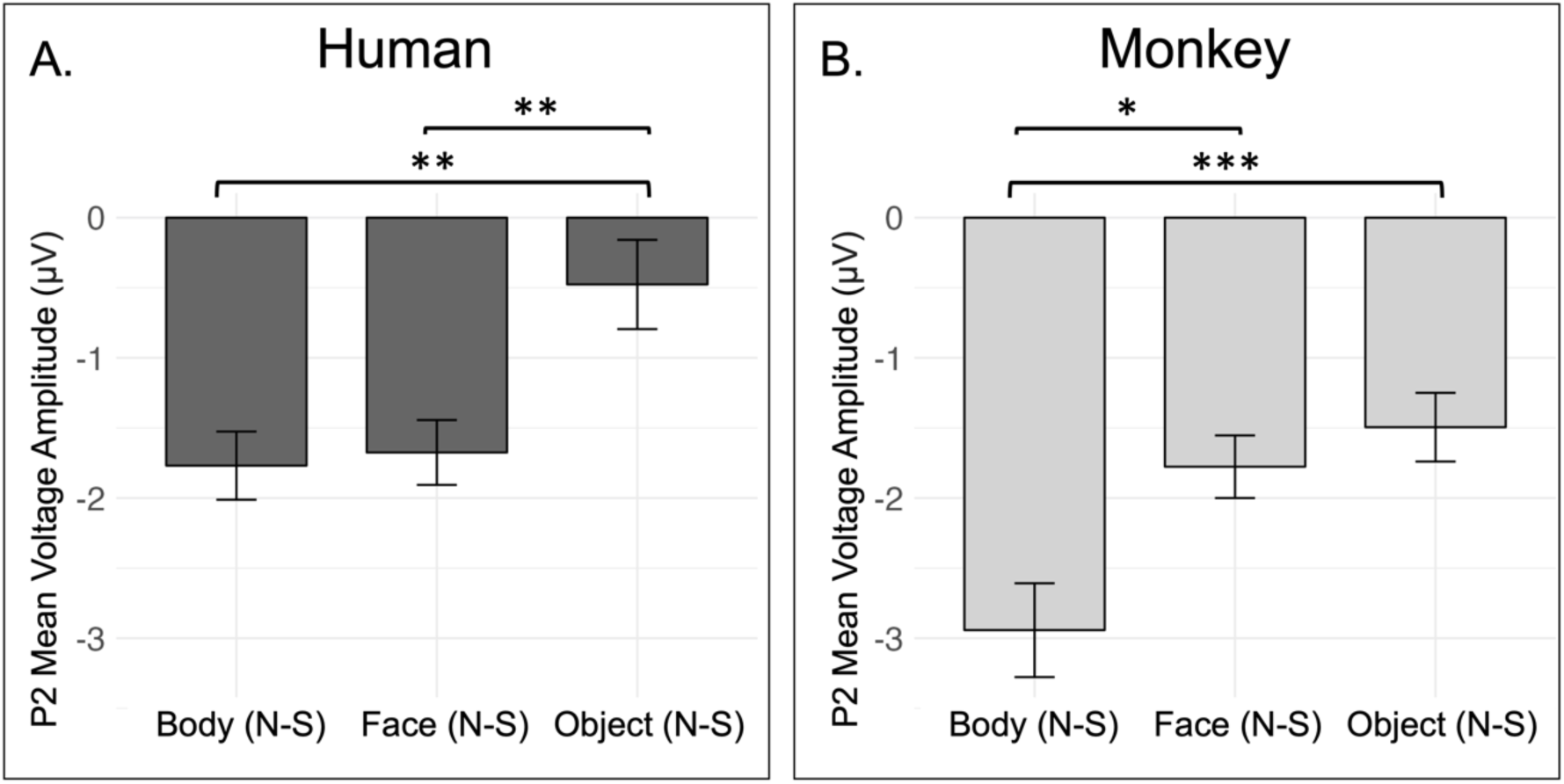
Means of differential voltage amplitude (Normal – Scramble) for component P2, shown separately for human categories (A) and monkey categories (B). As indicated by asterisks, a significant difference was observed between body and object categories for both human (*p*-adj = 0.003) and monkey (*p*-adj < 0.001). For human categories, a significant difference was observed between face and object (*p*-adj = 0.006). For monkey categories, a significant difference was observed between body and face (*p*-adj = 0.01).

Similar to these human-category results, analysis of monkey categories revealed a significant effect of category (*F*(2, 83) = 7.97, *p*-adj = 0.001, *η*^2^ = 0.16), driven by a significant difference between monkey body and monkey object categories (MD = 1.45, 95% CI [0.53, 2.36], *p*-adj < 0.001). Unlike the human-category results, there was a significant difference between monkey body and monkey face categories (MD = 1.17, 95% CI [0.24, 2.09], *p*-adj = 0.01) (Fig. 6B) and no significant difference between monkey face and monkey object categories (MD = 0.28, 95% CI [-0.64, 1.21], *p*-adj = 0.748).

## 4. Discussion

Our goal was to identify the temporal pattern of naturalistic body motion processing. Given the well-established EEG evidence for biological motion-evoked cortical responses, as well as recent fMRI evidence for naturalistic body-selective networks (see Introduction), we hypothesized that these naturalistic body motion processes would be reflected in ERP components. We further hypothesized that this effect would be observable with human body motion as well as monkey body motion.

To test our hypotheses, we recorded EEG from human subjects and employed a highly controlled stimulus set of naturalistic videos, marking the main methodological novelty of the present study. Videos depicted naturalistic body motion, face motion and object motion to disentangle body motion selectivity from visual features, such as luminance, contrast and non-background area, mosaic-scrambled versions of each stimulus were presented, and higher-level neural representations were derived from the normalization of Normal – Scramble ERP voltage. To further characterize these representations, animate stimuli depicted both human and monkey videos to probe the role of species in body motion processes.

In line with previous ERP literature on visual motion processing, video stimuli elicited strong N1 and P2 responses over bilateral temporo-parieto-occipital channels (Fig. 2), suggesting that our naturalistic videos indeed engaged global motion processes. More importantly, the normalized body, face and object motion responses during component P2 were all significantly different from zero (*p*-adj < 0.05) (Fig. 3), pointing to a higher-order motion process on a categorical level approximately 220 ms after video onset. The robustness of these higher-order motion processes is further supported by a sustained negativity following the strong, temporally concise response during component P2 and continuing until approximately 800 ms after video onset (Fig. 3).

The main finding of the present study is that the early stages of the aforementioned motion process are selective for moving stimuli of specific categories, in particular the body, which aligns with the hypothesis of naturalistic body motion-sensitive neural representations. This is shown by the statistical analysis of the higher-order, normalized ERP amplitudes, which showed the observed global motion effect was modulated by category for both components P2 and N2 (*p*-adj < 0.05), with bodies eliciting a stronger effect than objects (Fig. 4). Thus, at the stage of P2, the cortex may already distinguish between the global the case. In addition, as no statistically significant differences were observed between faces and objects, this effect cannot be merely attributed to differences in animacy.

These findings corroborate previous research on the temporal markers of biological PLD motion-specific processes at the stage of P2 (Buzzell, 2013; Pegna, 2015; Krakowski et al., 2011; Reid et al., 2008) and N2 (Hirai et al., 2003, 2005; Jokisch et al., 2005; White et al., 2014). Expanding on previous PLD investigations on body motion, the present study has, for the first time, shown that these motion-sensitive ERP markers are also distinguishable with naturalistic stimuli. While the aforementioned PLD studies localized the neural processes related to biological motion, they provided only limited insight to the higher-level, scramble-controlled representations of naturalistic movement. On a similar note, in line with recent fMRI research proposing two large-scale neural networks for naturalistic body motion in the lateral occipital cortex and right superior temporal sulcus in the human brain (Li et al., 2023), the present findings complement these spatial localizations with millisecond-precise temporal markers of the underlying neural processes.

Post-hoc investigations of component P2 suggested that the observed category effect on natural motion processing applies to motion of human bodies as well as monkey bodies (Fig. 5). Namely, significant differences were observed between body and object categories for both human and monkey videos (Fig. 6). These exploratory findings further point to a true, naturalistic body motion effect at the stage of P2. Yet, several questions remain unanswered; for example, future investigations should evaluate the specific cortical sources underlying the P2 body would allow directly correlating the present temporal markers with functional brain regions, possibly in right STS (Li et al., 2023) or a dorsal pathway for body motion information processing that has been proposed in computational models of body movement (Giese & Poggio, 2003).

As outlined above, the present study reports evidence for temporally distinct naturalistic body movement processes. This raises an interesting question about the relation between naturalistic human body processing and previous research revealing distinct processes for biomechanically plausible versus implausible movements (Candidi et al., 2008; Van Overwalle & Baetens, 2009). For example, a robust line of research points to distinct neural responses for biomechanical violations of hand movements (Longhi et al., 2015), finger movements (Costantini et al., 2005), and arm movements (Morita et al., 2012). Furthermore, recent models of EBA activity underscore the brain’s sensitivity to biomechanical plausibility (Marrazzo et al., 2024), paralleling the present findings of naturalistic body movement specific temporal processes. Future investigations are warranted to better understand this link, specifically the temporal profile of body movement-specific signal propagation to EBA and the putative influence of biomechanical violations.

In summary, traditional perspectives on category representation in the cortex and on body-specific processes are increasingly challenged by integrative perspectives highlighting the interaction of body processes with processes of biological movement (Jastorff & Orban, 2009), postural features (Marrazzo et al., 2023; Marrazzo et al., 2021; Poyo Solanas et al., 2020), action recognition (Goldberg et al., 2014; Shmuelof & Zohary, 2005) and social perception (Kret et al., 2011; of body processes. Moving forward, while mathematical models (Kumar et al., 2023) and theoretical frameworks (de Gelder & Poyo Solanas, 2021) have been proposed, a comprehensive model of body motion processing in the human brain is critical.

## 5. Data and code availability

The data and code that support the findings of this study are available on request from the corresponding author (Beatrice de Gelder), pending approval from the researcher’s local ethics committee and a formal data-sharing agreement.

## 6. Author contributions

Jane Chesley (Conceptualization, Data curation, Formal analysis, Investigation, Methodology, Visualization, Writing – original draft, Writing – review & editing, Project administration), Lars Riecke (Conceptualization, Methodology, Writing – review & editing, Supervision), Beatrice de Gelder (Conceptualization, Methodology, Writing – review & editing, Supervision, Funding acquisition).

